# Relative positioning of B and T cell epitopes drives immunodominance

**DOI:** 10.1101/2022.02.17.480954

**Authors:** Riccardo Biavasco, Marco De Giovanni

## Abstract

Humoral immunity is crucial for protection against invading pathogens. Broadly neutralizing antibodies (bnAbs) provide sterilizing immunity by targeting conserved regions of viral variants and represent the goal of most vaccination approaches. While antibodies can be selected to bind virtually any region of a given antigen, consistent induction of bnAbs in the context of influenza and HIV has been representing a major roadblock. Many possible explanations have been considered, however, none of the arguments proposed so far seems to fully recapitulate the observed counter-selection for broadly protective antibodies.

Antibodies can influence antigen presentation by enhancing the processing of CD4 epitopes adjacent to the binding region while suppressing the overlapping ones. We analyzed the relative positioning of dominant B and T cell epitopes in published antigens that elicit strong and poor humoral responses. In strong immunogenic antigens, regions bound by immunodominant antibodies are frequently adjacent to CD4 epitopes, potentially boosting their presentation. Conversely, poorly immunogenic regions targeted by bnAbs in HIV and influenza overlap with clusters of dominant CD4 epitopes, potentially conferring an intrinsic disadvantage for bnAb-bearing B cells in germinal centers.

Here we propose the theory of immunodominance relativity, according to which relative positioning of immunodominant B and CD4 epitopes within a given antigen drives immunodominance. Thus, we suggest that relative positioning of B-T epitopes may be one additional mechanism that cooperates with other previously described processes to influence immunodominance. If demonstrated, this theory can improve the current understanding of immunodominance, provide a novel explanation on HIV and influenza escape from humoral responses, and pave the way for new rational design of universal vaccines.

## Introduction

Antibodies are a fundamental component of the human immunological defense, and one of their most important functions is to confer protection against viruses and exogenous microorganisms upon primary infection or vaccination^1–3^. Antibodies that are able to provide sterilizing immunity, by preventing pathogen entry and interaction with target cells, are commonly called ‘neutralizing antibodies’ (nAbs)^1^. nAbs are produced by B cells that have been selected in the host germinal centers (GC)^3–5^. Upon antigen encounter, antigen-specific B cells start proliferating, interact with cognate CD4 T cells, and then migrate to the center of B follicles where they establish the GC microstructure^3,4,6–8^. GCs are the microanatomical niches where B cell clones mature, receive survival signals by cognate CD4 T cells, mutate their B-cell receptors (BCRs) and are selected to produce high-affinity nAbs^3,5,9^.

The repertoire of naïve B cells and BCRs is extremely diverse thanks to BCR rearrangement during B cell development, resulting in the potential to produce antibodies against virtually any epitope of a given antigen^2,10,11^. Despite this potential, epitope specificities are not equally targeted by humoral responses, with the most frequently targeted epitopes defined as immunodominant^3,12^. Following immunization, the higher prevalence of specific B and T cell clones, which expand at the expense of other epitope-specific cells, is referred to as immunodominance^3,12^. The combination of germline B cell precursor frequency, antigen accessibility, affinity/avidity and CD4 T cell help influence the Darwinian selection process of B cell clones in GCs and largely impact on epitope immunodominance^1,3^.

The goal of most vaccines is to induce vigorous and long-term production of nAbs with the ability to prevent any future infection. However, some viruses deploy diverse strategies to escape antibody neutralization, among others, the hypermutation of viral antigens (e.g. HIV, influenza)^1,3,13^. Antibodies that bind to conserved viral epitopes and are able to neutralize different viral mutants and strains are referred to as ‘broadly neutralizing antibodies’ (bnAbs)^1^. Conserved viral epitopes often represent the ‘Achilles heel’ of mutating viruses, as they cannot be mutated without altering important steps in the viral life cycle^1,14,15^. While the induction of bnAbs represents the major goal of most vaccination approaches, none of the strategies tested so far has successfully and consistently induced anti-HIV and anti-influenza bnAbs^13,15,16^. Indeed, immunodominance of B cell clones specific for variable and/or non-broadly neutralizing viral epitopes has recently proven to be a major obstacle in vaccine design, with enhanced production of non-neutralizing antibodies at the expenses of the bnAbs^3,12,15,16^.

Poor or defective bnAbs induction following vaccination in the context of HIV and influenza has been extensively studied. Potential explanations for defective bnAbs production (e.g. anti-gp120 CD4-binding site for HIV, anti-stem of the hemagglutinin (HA) for influenza) have been proposed: poor epitope accessibility^3,17,18^, high mutational load required to generate bnAbs^1^, low neutralizing B cell precursor frequency^19^ and HLA-II polymorphisms^20,21^. Nevertheless, none of these arguments can fully recapitulate the apparent counter-selection for bnAbs, with recent experimental evidence arguing against such theories.

Around 30 years ago, antibody binding was shown to influence antigen processing and presentation, with CD4 T cell epitopes either inhibited or boosted based on their relative positioning to the antibody epitope^22–24^. Here, we propose the theory of immunodominance relativity, according to which the relative positioning of B and T cell epitopes within a given antigen drives immunodominance. During a global pandemic caused by a newly emerged coronavirus, understanding the molecular bases of immunodominance has become of paramount importance, particularly to guide the rationale design of future universal vaccines.

## Historical viral escape theories

Over the past decades, several hypotheses have been proposed to explain the molecular mechanisms underlying the inconsistent induction of bnAbs, especially in the context of HIV and influenza vaccination studies.

### Antigen variability and epitope accessibility

Variation of immunodominant epitopes due to accumulation of random mutations in the viral genome represents one of the most successful strategies for viruses to escape the host immune system^1,13^. This strategy is particularly relevant for RNA viruses, which rely on more error-prone molecular machinery when duplicating their genome^25^. However, HIV and influenza envelope proteins (gp120 and HA, respectively) contain 3-dimensional structures that are not permissive to high mutational load, as this would result in impairment of essential steps during the viral life cycle, such as binding to target receptors, membrane fusion and internalization^1,25,26^. Indeed, conserved regions of these antigens, namely the CD4-binding domain of gp120 and the stem of HA, contain the epitopes targeted by the most potent bnAbs described so far against these two viruses, 3BNC117^27^ and MEDI8852^28^, respectively. Nevertheless, antibody responses targeting the CD4-binding domain and the stem of HA are very rare in the general population.

Epitope accessibility has long been regarded as a fundamental requirement to effectively mount humoral responses^1,14,18,29–32^. First-generation anti-HIV bnAbs targeting the CD4-binding site showed limited neutralization breadth and/or potency, a finding thought to be related to poor accessibility of these epitopes in HIV-1 primary isolates^33^. Recently, however, a renewed experimental effort has led to the isolation of several new anti-CD4 binding site human monoclonal antibodies (VCR01-like) characterized by greater breadth and potency^19,34,35^. Moreover, the CD4-binding site is accessible for binding to human CD4 molecules^14^, an essential step during viral entry into target cells, and VRC01-like Abs can be generated in macaques following vaccination^36^, indicating that this region can be immunogenic *in vivo*. In the context of influenza HA, steric hindering from the head domain has been considered to prevent Ab binding to the stem domain^1,15^. Nevertheless, further increasing the accessibility of the stem with head-less HA immunogens didn’t result in potent bnAbs induction^1,37^, overall suggesting that low accessibility of conserved viral epitopes does not fully explain mechanisms for subdominant bnAbs induction. More recently, bnAbs were shown to efficiently bind to cryptic epitopes in different microbial antigens, such as coronavirus^38^, ebolavirus^39^, and plasmodium^40^, conferring neutralization across different subspecies of those pathogen families.

### B-cell precursor frequency, somatic hypermutation and HLA2 polymorphisms

Frequency of germline B cell precursors shapes the humoral response to immunogenic antigens by impacting GC occupancy^1,3^. Low B cell precursor frequency has been proposed as one potential obstacle in mounting effective bnAb responses following vaccination. Despite being relatively rare in the repertoire, VRC01-like cells can be activated by high-affinity stimulation in mouse models that recapitulate human precursor frequencies^3,18^. VRC01-like precursors are also present in 96% of humans^19^, giving hope that a bnAb-like HIV vaccine is indeed possible. Furthermore, germline residues of the VH1-69 alleles, which account for 2-6% of B cell repertoire, are known to mainly mediate recognition of influenza group 1 HA-stem^1,41^. This extraordinary high number of putative anti-stem B cell precursors correlates, however, with subdominant VH1-69 response that is generated only in some individuals.

The high mutational load required to develop bnAbs can also represent a major challenge^1^. Indeed, anti-HIV VRC01-like antibodies require multiple rounds of somatic hypermutation to be generated and are frequently characterized by extensive nucleotide insertions and substitutions^1,3^.. Nonetheless, it was recently shown that a single proline-to-alanine mutation in HCDR2 is sufficient to confer high affinity binding to influenza HA-stem^41^, consistently with rapid affinity maturation process. Despite the low number of mutations required, anti-influenza bnAbs are rare and apparently counter-selected, suggesting that high mutational load and complex antibody selection processes do not fully recapitulate defective bnAbs induction, at least in the context of influenza.

A variety of host genetic factors can influence the outcome of viral infections, most notably polymorphisms within the HLA class I and II loci^20,42–44^. HLA class II molecules are essential in the development of adaptive immune responses, as they present antigens to CD4 T cells and contribute to the regulation of B and T cell interactions within GCs. HLA DRB1*13-DQB1*06 was associated with a trend toward increased duration of AIDS-free time in HIV patients treated with anti-retroviral therapy^43^. In addition, inheritance of DRB1*13 alleles has been associated with long-term survival among children with vertically transmitted HIV-1 infection^20^ and higher IFNγ production by Th1 cells^21^. Nevertheless, anti-CD4 binding site (and gp41-MPER) bnAb responses were detected in very few individuals independently of HLA class II haplotypes^45^, suggesting that HLA class II polymorphisms are not likely to explain the defective anti-viral bnAbs production following vaccination.

### BCR and antibody modulation of antigen presentation

Antigen processing and presentation was initially investigated using macrophages or dendritic cells as a model of antigen-presenting cells (APCs)^46^. In such cases, the APCs do not possess specific antigen receptors and antigen internalization is mainly restricted to receptor-free endocytosis. In contrast, antigen uptake by antigen-specific B lymphocytes is mediated by the BCR, which mediates efficient antigen presentation at lower concentrations than those required by non-specific B cells^23,47^. Antibodies can alter the conformation and stability of target antigens^48^ and protect epitopes from proteolytic processing^49^. In particular, antibody-antigen complexes can resist the lysosomal acidic pH^22^, therefore influencing the antigen fragmentation by proteases, which can initiate while the antigen is still bound by antibodies^50^.

Pioneer work by *Berzofsky* and *Celada* indicated that antibody binding could differentially boost antigen presentation to some T cell clones, either through modulation of antigen uptake or proteolytic processing^51–54^. Accordingly, it was hypothesized that the antibody specificity shapes the initial pattern of antigen fragmentation, supporting the existence of T and B cell preferential pairing^51,55^ and reciprocity circuits^51^. Few years later, elegant work from *P*.*D. Simitsek et al* demonstrated that antibody binding to antigens can modulate their processing by enhancing or suppressing HLA class II presentation of different CD4 T cell determinants^56^. Strikingly, a single bound antibody (11.3) or its Fab fragment was shown to simultaneously boost the presentation of one CD4 T cell epitope (1273-1284aa), while suppressing the *in vitro* presentation of other determinants (1174-1189aa, **Fig. S1**)^56^. Both tetanus epitopes that are modulated by BCR/antibody binding were shown to fall within the 11.3 Ab footprint region (i.e. the region of the antigen that is at least partially protected from lysosomal proteolytic cleavage). The suppressed epitope (1174-1189aa) is sterically hindered to bind to HLA class II molecules upon interaction with 11.3 Ab, likely due to reduced proteolysis in the lysosomal and inaccessibility to HLA binding^56^. On the contrary, the boosted epitope (1273-1284aa) was likely protected from excessive cleavage, stabilized, and made more accessible for the binding to HLA class II molecules by the interaction with the 11.3 Ab. Processing of epitopes located far from the BCR/antibody binding region (947-967aa) was instead maintained unaltered. The influence of the antibody specificity on antigen degradation was further analyzed in clones of tetanus-specific B cells with different epitope specificities^50,57^, and was confirmed using other model antigens, such as f3-galactosidase^52^ and myoglobin^54^.

## Results

### Strong inducers of neutralizing humoral responses

Given the impact of BCR and antibody binding on antigen processing, we postulated that the relative positioning of B and T cell epitopes shapes immunodominance. To test this hypothesis in the absence of studies that map B and T immunodominant epitopes within single patients, we analyzed the 3-D positioning of published B and T cell epitopes in immunogenic antigens known to elicit strong and long-lasting neutralizing humoral responses, such as measles HA, diphtheria toxoid, vesicular stomatitis virus glycoprotein (VSV-GP) and SARS-CoV-2 spike protein. To restrict our focus on highly immunodominant and HLA-independent regions, we gathered published data for B and T cell epitopes and selected determinants found in multiple patients or host species (mice, macaques and humans).

Across measles HA and diphtheria toxoid, immunodominant CD4 epitopes are scattered throughout the protein sequences and often adjacent to immunogenic and neutralizing B cell epitopes (e.g. Measles HA: T epitopes 321-350^58^, B epitopes 309-319, 380-400^59^; Diphtheria toxoid: T epitopes 271-290, 321-340, 411-450^60^, B epitopes 247-260, 395-403, 477-483, 508-527^61^, **Fig. 1a, b; Table S1**,**2**). In the diphtheria toxoid (PDB 1MDT), 5 out of 18 immunodominant B regions (255-260, 381-394, 395-403, 452-458 and 465-475aa) are located next to immunogenic CD4 epitopes (271-290, 351-371, 411-430, 431-450aa), possibly in the ideal position to be boosted upon BCR binding (**Fig. 1b**). Interestingly, only 2 out of 18 immunodominant B cell epitopes (351-355, 409-420aa)^61,62^ partially overlap with immunodominant CD4 determinants (351-370, 411-430aa)^60^, however they are also adjacent to other CD4 determinants (331-350, 421-440aa) (**Fig 1b** and **Table S2**). In VSV-GP (PDB 4YDI), a neutralizing B cell epitope (382-400aa)^63^ is located next to 2 immunodominant CD4 epitopes (338-368aa)^64^ and does not overlap with any CD4 epitope. Protective antibodies targeting the spike protein of SARS-CoV-2 are induced in the majority of infected or vaccinated patients^65–67^. In agreement with previous observations on immunogenic antigens, 4 out of 8 B cell immunodominant epitopes (209-226, 721-733, 769-786, 809-826aa)^68,69^ are adjacent to immunodominant CD4 determinants (166-180, 751-765, 866-880aa)^70^ in SARS-CoV-2 spike protein, possibly inducing antigen presentation boost upon BCR binding (**Fig. 2** and **Table S4**). In parallel with other highly immunogenic antigens, none of the immunodominant B regions are found to be overlapping with dominant CD4 epitopes. Despite broad immunogenicity, the spillovers of β-coronaviruses in humans and the emergence of SARS-CoV-2 variants highlight the need for broader anti-coronavirus humoral protection. Recently, a highly conserved B epitope in the stem-helix of SARS-CoV-2 spike (1148-1156aa) was shown to be target of several bnAbs^66^, thus representing a potential target for broad humoral protection. Higher frequencies of anti-stem helix-specific Abs were observed for vaccinated individuals who were previously infected^66^, indicating this region is immunogenic in humans. However, these antibodies are found at much lower frequency in individuals previously infected with SARS-CoV-2 or those who had received two doses of mRNA vaccines, indicating that humoral responses targeting stem-helix are usually rare. Interestingly, this highly conserved stem-helix B epitope (1148-1156aa) is not located near any immunodominant CD4 determinant, where antigen presentation boost upon BCR binding is unlikely to take place. This observation can explain the counter-selection of anti-stem-helix bnAbs in favor of more immunogenic but less cross-reactive anti-spike B cell specificities, which are adjacent to immunodominant CD4 determinants and might take advantage of enhanced antigen presentation. Of interest, other neutralizing and dominant B epitopes were described to locate within the RBD spike domain, distant from dominant CD4 determinants^71,72^

**Fig. 1.**
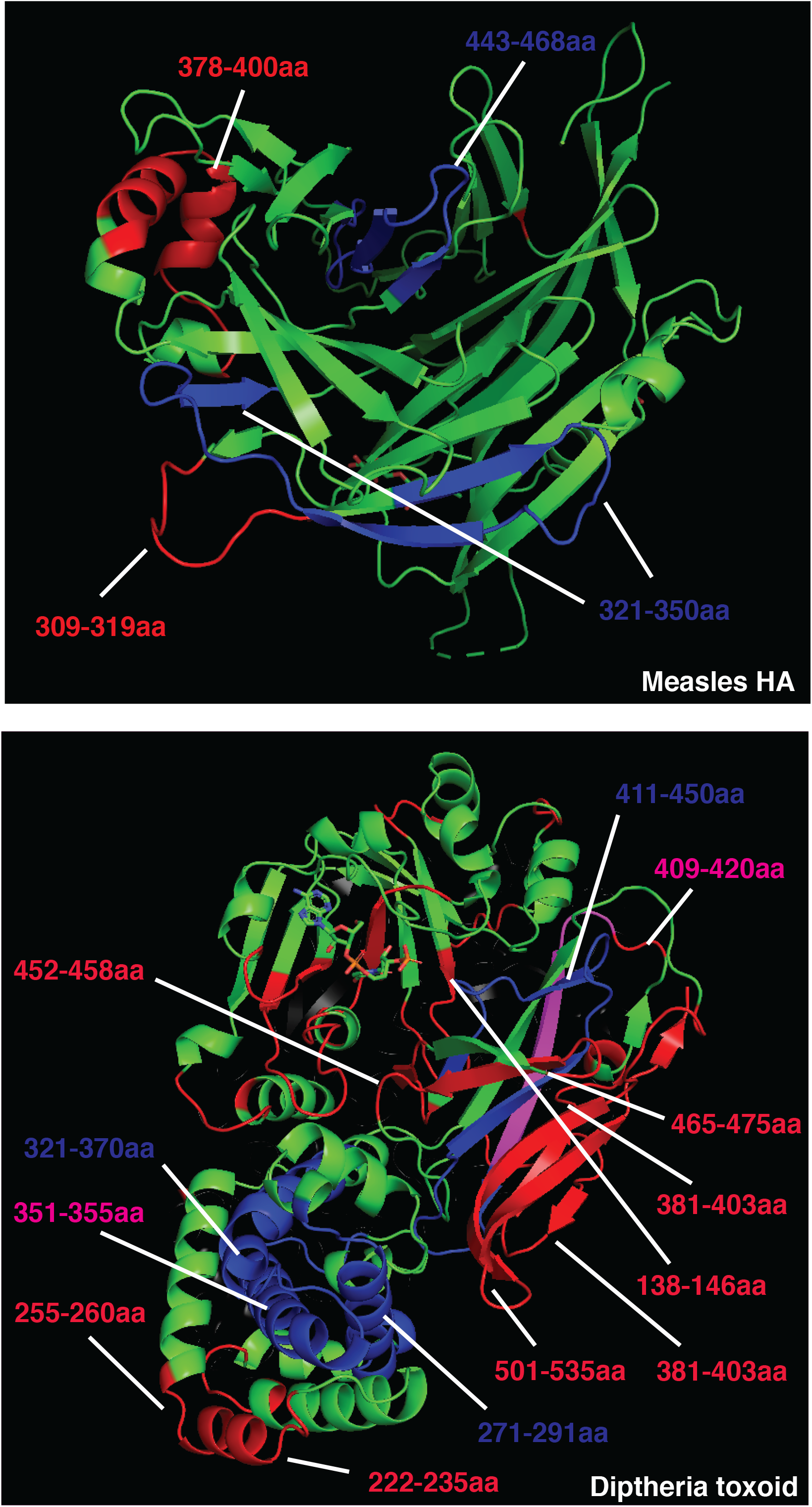
Crystal structure of Measles hemagglutinin (**PDB:2ZB6, a**) or Diptheria toxoid (**PDB:1MDT, b**). Immunodominant B cell epitopes are depicted in red, while dominant CD4 determinants are highlighted in blue.

**Fig. 2.**
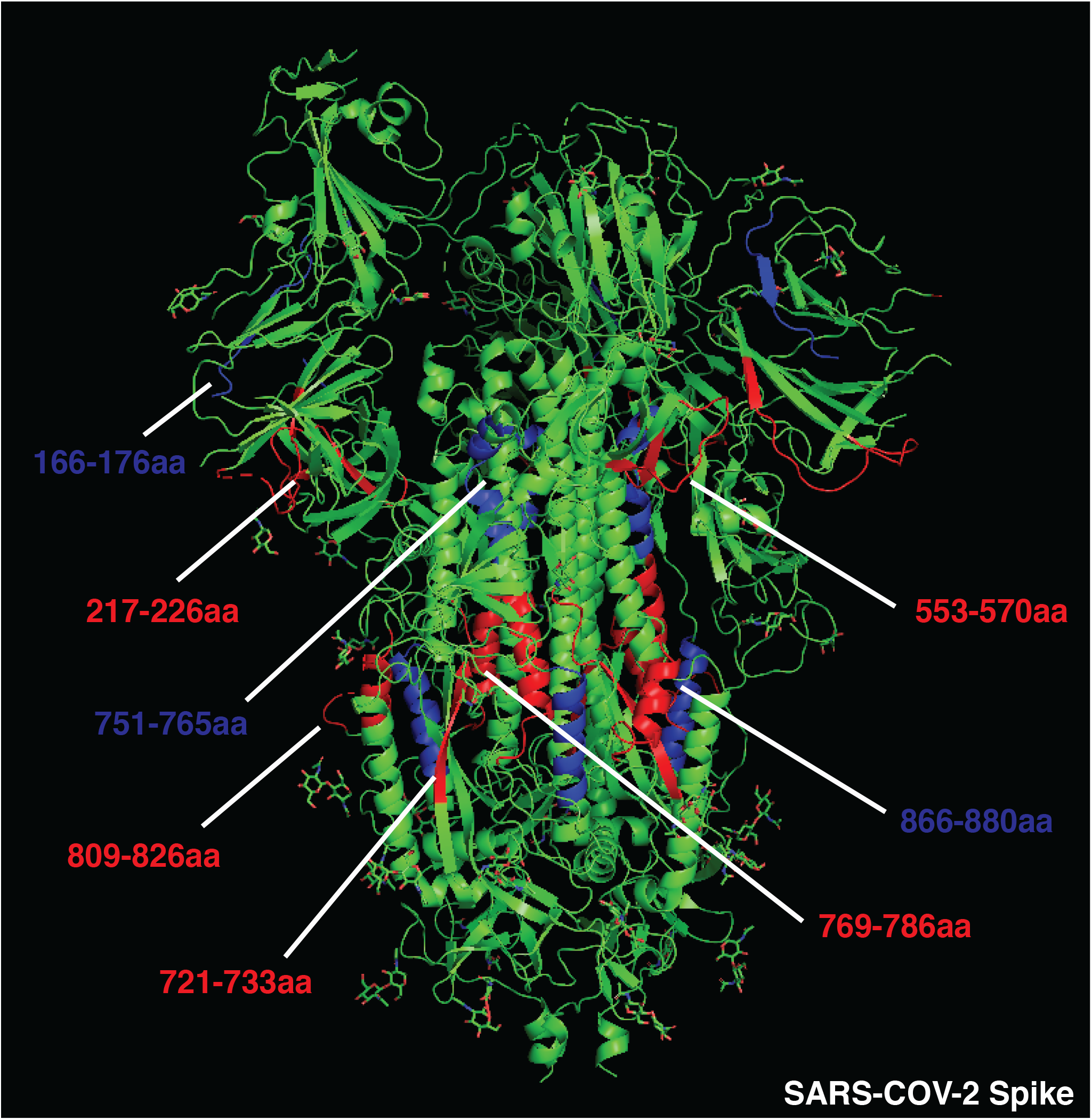
Crystal structure of SARS-COV-2 spike protein (**PDB:6VSB**). Immunodominant B cell epitopes are depicted in red while, dominant CD4 determinants are highlighted in blue.

Overall, these data suggest that immunodominant and neutralizing B cell epitopes are mostly not overlapping with and often adjacent to dominant CD4 T cell epitopes, increasing the chances of antigen presentation boost rather than suppression upon BCR binding. Additionally, antigens that are associated with rapid nAb response contain immunogenic CD4 determinants scattered in several portions of the viral proteins. Scattered CD4 T cell epitopes increase the likelihood for dominant B cell clones targeting multiple regions of the antigen to emerge, facilitating the neutralizing humoral response.

### Poor inducers of neutralizing humoral responses

In addition, we postulated that clustering of dominant CD4 T cell epitopes could reduce the immunogenicity of specific antigen portions, suppressing T cell help to B cells targeting those regions. To test this hypothesis, we modeled the relative positioning of B and T cell epitopes in well-studied antigens that efficiently escape broad Ab neutralization such as HIV gp120 and influenza HA. To minimize the effects of viral antigen variability and HLA polymorphisms, we focused on experimentally validated and highly immunodominant CD4 determinants presented by multiple HLAs (see methods) and conserved B cell epitopes targeted by bnAbs.

The CD4-binding site in HIV gp120 represents, among others, one of the most promising targets to achieve broad anti-HIV protection. Indeed, second-generation anti-CD4 binding site antibodies (e.g. 3BNC117) broadly neutralize HIV-1 primary isolates and suppress infection upon intravenous injection in chronically infected patients, representing a potent clinical tool. Intriguingly, immunodominant CD4 epitopes are not scattered throughout the whole gp120 protein, rather they are clustered in the outer domain^73^ (**Fig. 3a**). In addition, combining rules for HLA class II binding of predicted epitopes to well-conserved sequences substantially improves the prediction of immunodominant CD4 epitopes^74^, as epitopes included in conserved sequences are more likely to become immunodominant thanks to a higher frequency across different viral strains. Altogether, these observations suggest that HIV might have evolved to host a cluster of immunodominant CD4 T cell epitopes within the CD4 binding site, a highly conserved region of the gp120 outer domain and the target of the most promising bnAbs (**Fig. 3a**). To confirm this observation, we selected validated immunodominant CD4 epitopes published by different research groups, focusing on those shared by multiple HLA class II haplotypes (immunodominant in at least two different mouse strains, macaques and/or patients). Highly immunodominant CD4 epitopes are localized within 3 main regions in gp120 sequence: 300-368, 400-449 and 480-508aa^73–77^ (**Fig. 3** and **Table S5**), confirming the clustering of immunodominant epitopes within CD4 binding site in the outer domain. Crystal structures of HIV gp120 and 3BNC117 bnAb (PDB 4jpv) are shown in **Fig. 3b**. The neutralizing epitope recognized by 3BNC117 is located within the CD4-binding site and overlaps with immunodominant CD4 T cell determinants (300-368, 400-449 and 480-508aa). Strikingly, 62% of the amino acids essential for 3BNC117 binding (**Fig. 3b** and **Table S5**) are located within the highly immunodominant CD4 T cell epitopes. High overlap of B and T cell epitopes may result in suppression of antigen presentation and intrinsic disadvantage of B cells displaying 3BNC117^19^-like BCRs. In support of this hypothesis, high-affinity anti CD4-binding site Abs can inhibit gp120-specific CD4 T cell proliferation if added during APC-Ag pulsing^78^ by suppressing gp120 processing and preventing HLA class II antigen presentation^79^. Moreover, recent experimental evidence suggests that potent help is required to stimulate and expand rare precursors of anti-gp120 bnAbs *in vivo*, highlighting the importance of T cell help during this process^80^.

**Fig. 3.**
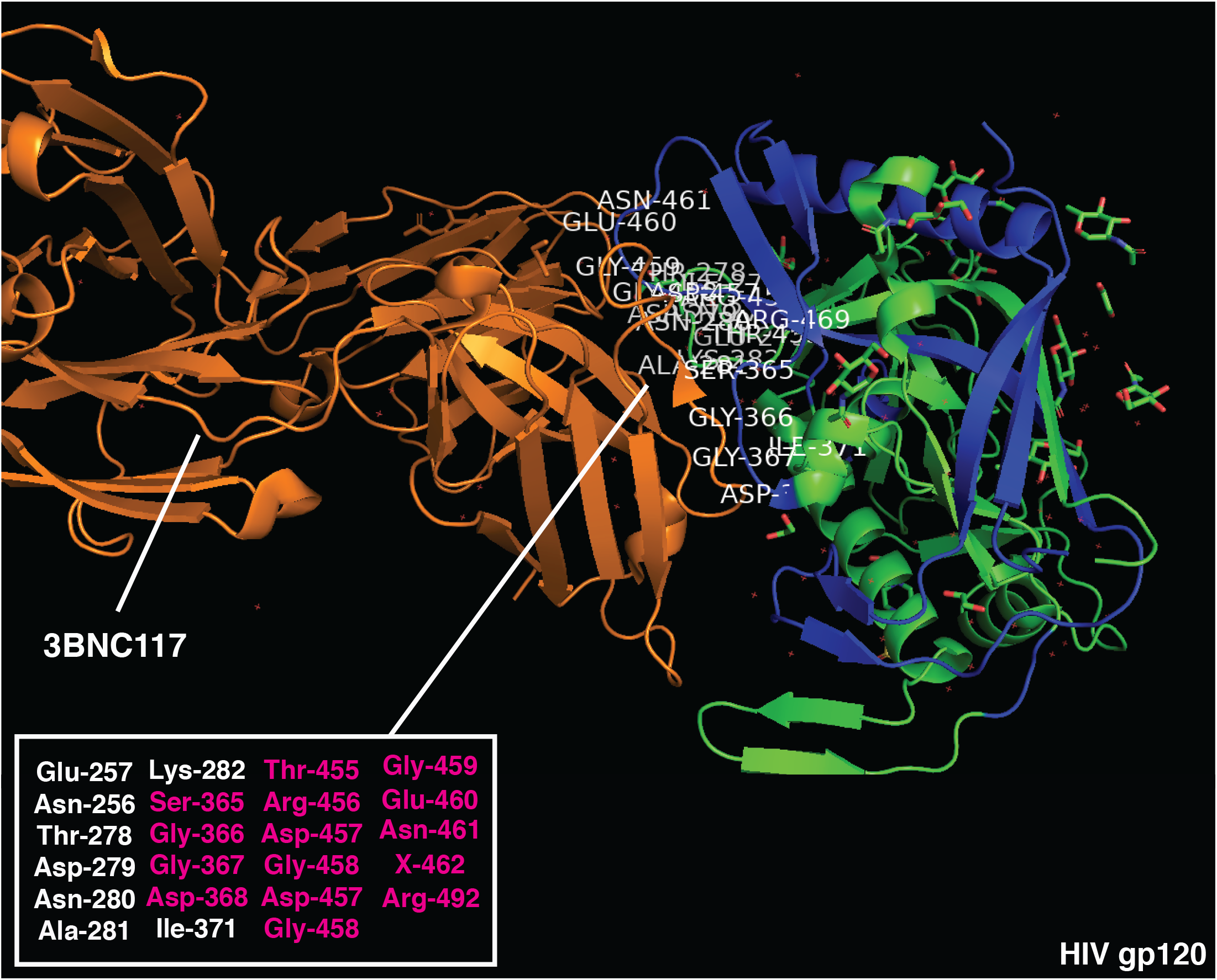
Crystal structure of HIV gp120 monomer and 3BNC117 bnAb (**PDB:4JPV**). Immunodominant CD4 epitopes are highlighted in blue. Amino acids essential for 3BNC117-binding are listed (bottom left) and those overlapping with CD4 dominant regions are highlighted in red.

Similarly, immunodominant CD4 T cell epitopes (400-430aa)^17^ largely overlap with neutralizing B cell regions in the influenza HA-stem (PDB 5JVR, **Fig. 4** and **Table S6**), with 50% essential ammino acids for MEDI8852 binding to HA-stem located within the highly immunogenic CD4 T regions (401-430aa). In support of this observation, anti-stem (but not anti-head) antibodies have recently been shown to specifically inhibit presentation of immunodominant T cell epitopes located within the HA-stem^17^. Influenza HA-head is the target of most strain-specific nAbs that commonly lack broadly neutralizing activity. Anti-HA-head B cell clones largely dominate the humoral responses, allowing for strain-specific nAbs to emerge in virtually every infected host. Of note, influenza HA-head also contains immunodominant CD4 epitopes (centered around 215 and 265 aa^81^), which are mostly adjacent to immunodominant B cell regions (**Fig. 4 and Fig. S3**. This relative positioning recapitulates the structure observed in highly immunogenic antigens, likely leading to enhanced antigen presentation upon BCR binding. On the contrary, the surface of the HA-stem region is constituted of immunodominant CD4 T cell epitopes, maximizing the likelihood of suppressing any B cell clone targeting this conserved protein region.

**Fig. 4.**
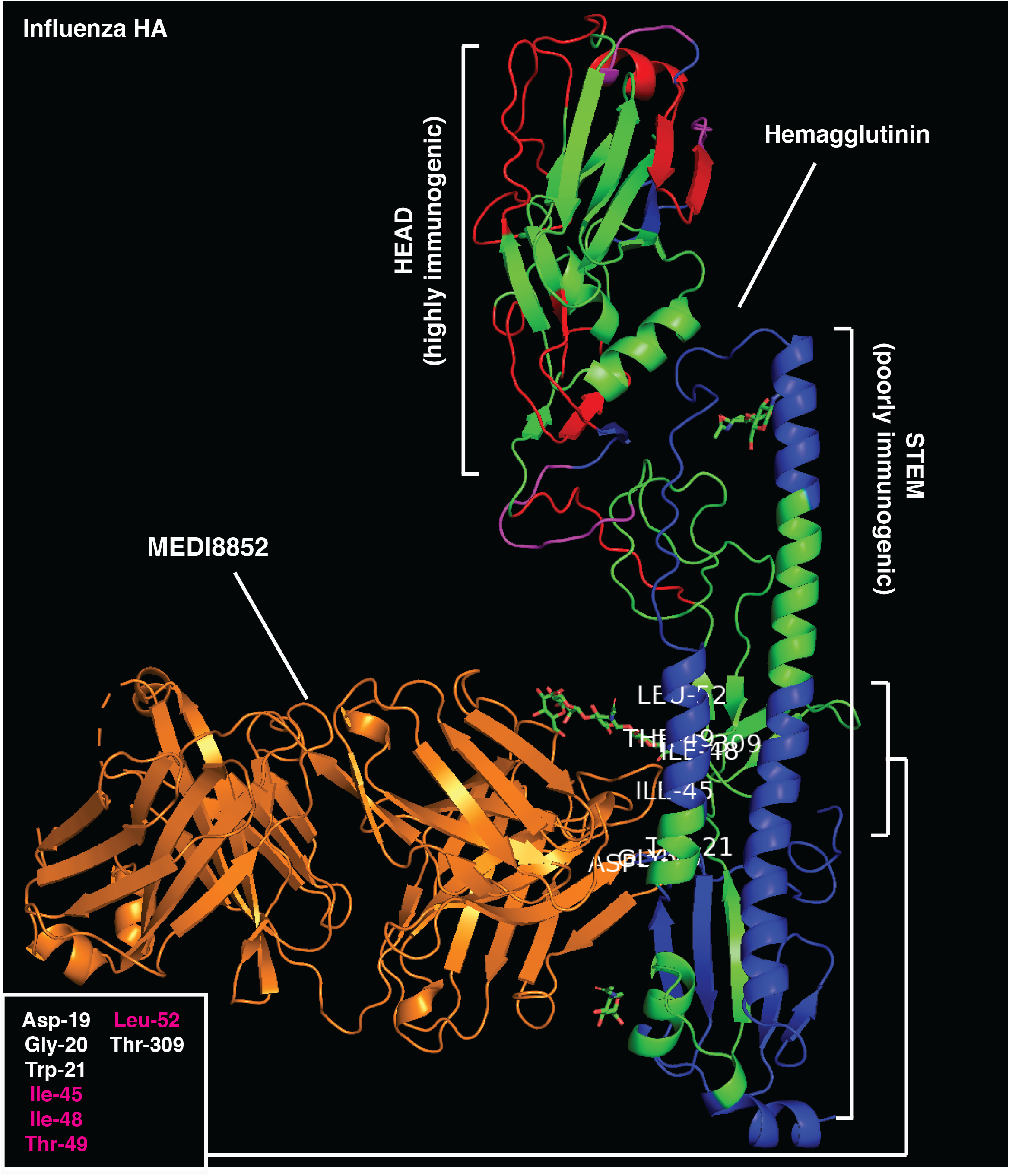
Crystal structure of influenza hemagglutinin and MEDI8852 bnAb (**PDB:5JW3**). Immunodominant B cell epitopes are depicted in red while, dominant CD4 determinants are highlighted in blue. Amino acids essential for MEDI8852-binding are listed (bottom left) and those overlapping with CD4 dominant regions are highlighted in red.

To quantify the observed differences in the relative positioning of B and T cell epitopes, we grouped together antigens that are strong or poor inducers of long-lasting humoral responses and measured the frequency of B epitopes that are distant (>15aa), adjacent (<15aa), or overlapping (0 aa) with CD4 T cell epitopes. In support of our hypothesis, the relative positioning of B and T cell epitopes is highly associated with the immunogenicity of the antigens analyzed (*X*^2^ (2,n=66)=23.05, p<0.0001, **Fig. 5a, S3** and **Table S7**), with poor immunogens showing increased frequency of B epitopes overlapping with immunodominant CD4 determinants compared to strong nAb inducers (58.6% vs 8.1%, respectively).

**Fig. 5.**
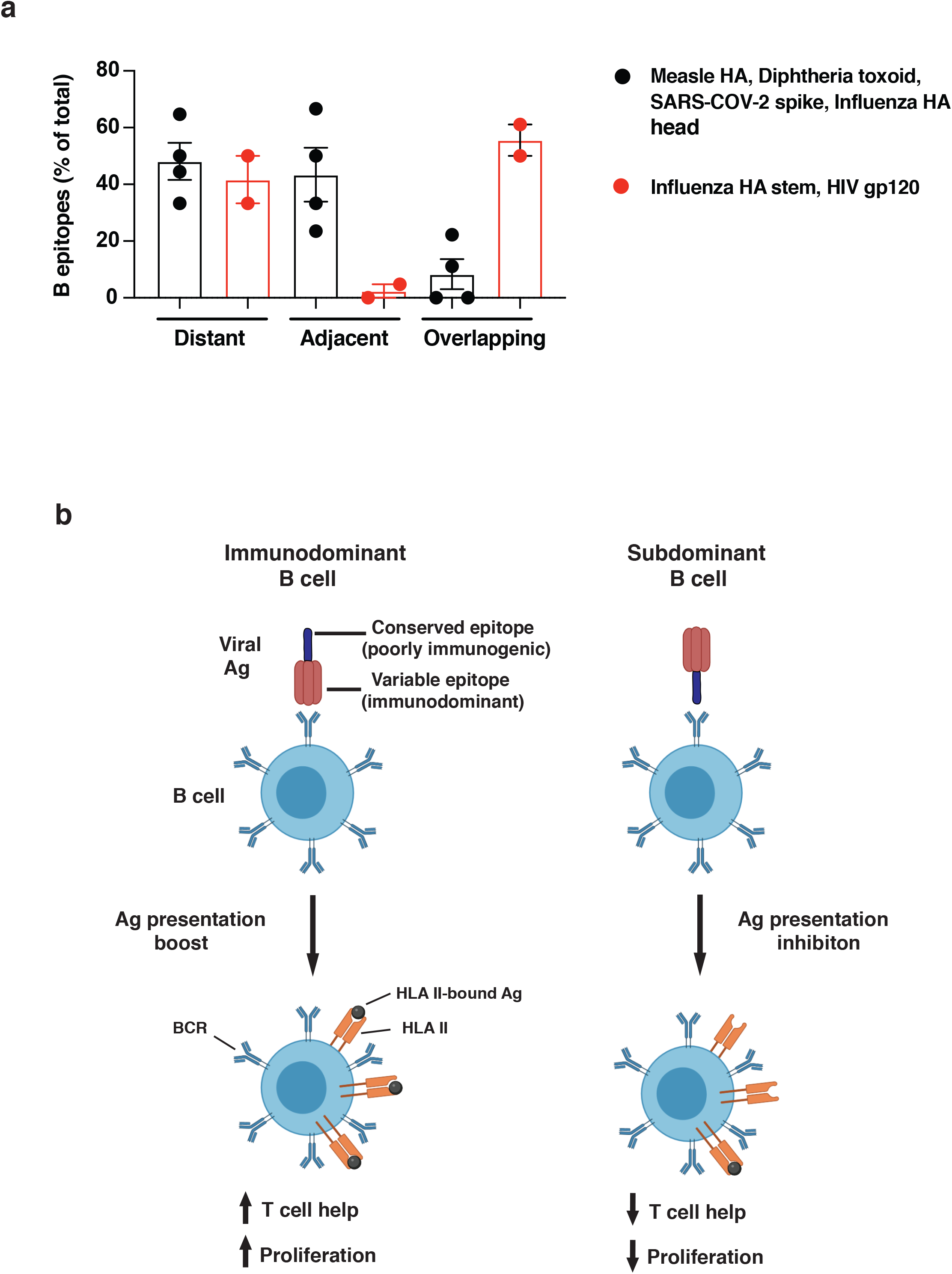
**a**. Quantification of dominant B cell epitopes classified as distant (>15aa), adjacent to (<15aa) or overlapping with (<0aa) immunodominant CD4 determinants within the same antigen. Good nAb inducers (measles HA, diptheria toxoid, SARS-COV-2 spike and influenza HA head, black dots) and poor nAb inducers (influenza HA stem and HIV gp120, red dots) are compared. Kruskal-Wallis test was applied. ^*^P <0.05. **b**. Model figure showing on left, immunodominant B cell (left) recognizing immunogenic viral regions and boosting antigen presentation upon BCR binding; right, subdominant B cell clone binding conserved / poorly immunogenic viral regions and inhibiting presentation of overlapping CD4 epitopes.

Altogether, this analysis supports the immunodominance relativity model, according to which the relative positioning of B and T cell epitopes within antigens drives B cell immunodominance. Epitopes targeted by bnAbs are often overlapping with highly immunodominant CD4 T cell epitopes within viral antigens that escape Ab neutralization (**Fig. 5b**). Epitope overlap may result in suppression of antigen presentation, limiting T cell help and introducing an intrinsic disadvantage for B cell clones displaying bnAb-like BCRs. To maximize this effect, immunodominant CD4 T cell epitopes are mostly scattered across the viral antigens that induce efficient Ab neutralization, whereas they tend to cluster within poorly immunogenic regions in antigens that escape humoral responses. In conclusion, the immunodominance relativity model offers an innovative explanation on HIV and influenza escape from long-lasting immunity and on the molecular basis of antibody selection and maturation.

## Discussion

Antibodies can potentially target any epitope of a given antigen, thanks to extremely high variability in the repertoire of B cell clones. Despite this potential, epitope specificities are not equally targeted by humoral responses, with the most frequently targeted epitopes defined as immunodominant. Viruses have evolved different strategies to escape Ab neutralization, among others hypermutation of viral antigens. Antibodies able to neutralize multiple viral variants are defined as broadly neutralizing and they represent the ultimate target of most vaccination strategies. BnAbs represent a fundamental tool to mount effective protection against highly mutating viruses such as influenza and HIV. Despite prolonged efforts to induce such humoral responses upon vaccination, bnAbs are generally highly subdominant when compared to other strain-specific antibodies.

Understanding the rules defining immunodominance is of paramount importance to improve the design of future vaccination strategies. Over the past decades, several hypotheses have been suggested to explain the molecular mechanisms underlying the inconsistent induction of bnAbs, including antigen mutation, epitope accessibility, high BCR-mutational load required, B cell precursor frequencies and HLA class II polymorphisms. Nevertheless, none of these hypotheses fully recapitulates the counter-selection of bnAbs, especially in the context of isolated antigens in vaccination studies. Pioneering work by *Berzofsky* and *Celada* demonstrated that BCR or antibody binding to antigens could boost or inhibit the presentation of specific CD4 epitopes based on their relative positioning. Indeed, antibodies can both sterically hinder and inhibit the presentation of CD4 T cell epitopes located within the Ab-bound region or stabilize adjacent epitopes to facilitate their mounting on HLA molecules, thus enhancing their presentation.

In this work, we propose the theory of immunodominance relativity, according to which the relative positioning of B and T cell epitopes within an antigen shapes immunodominance. Indeed, we found that subdominant conserved regions targeted by bnAbs (e.g. CD4-binding site of HIV gp120 or the stem of influenza HA) are often overlapping with clusters of highly immunodominant CD4 epitopes. In support of this observation, anti-CD4 binding site and anti-HA-stem Abs inhibit antigen presentation upon binding, reinforcing the idea that overlapping B and T cell epitopes can lead to inhibition of antigen presentation. Our analysis suggests that bnAb B cell precursors specific for conserved regions of HIV and influenza inhibit the presentation of immunodominant CD4 determinants upon BCR binding, resulting in poor T cell help during the GC reaction and consequent counter-selection in favor of other dominant B cell clones. On the contrary, non-neutralizing or strain-specific immunodominant B cell precursors may boost the presentation of adjacent immunodominant CD4 epitopes, resulting in increased T cell help and selective advantage during GC reactions. It is worth highlighting that the mechanism we are proposing here might have different impacts on B cell immunodominance depending on the B epitopes analyzed. Indeed, specific bnAb-targeted B cell determinants may be subdominant as a function of other immunosuppressive mechanisms, which are extensively discussed above. Interestingly, potent anti-HIV bnAb antibody can be eventually isolated from infected patients years after the infection. A possible explanation for this event in light of our theory is that extensive T-B crosstalk and BCR mutational lode in GCs over the years might somehow overcome defects in antigen presentation by bnAb-bearing B cell clones. We would also like to point out that a conformational (rather than linear) epitope analysis might impact on epitope interactions and will be of high interest to obtain in this context. Nevertheless, our analysis likely comprises both linear and conformational epitopes, as immunodominant B regions provided likely contain both linear and structural determinants.

This working hypothesis currently lacks formal experimental demonstration; however, we built the model gathering data from a multitude of independent publications from the last 4 decades and found supportive experimental evidence published by different groups. Finally, we propose that relative positioning of B-T epitopes may be one additional mechanism that cooperates with the other above-mentioned processes to influence immunodominance. If demonstrated, this theory can improve the understanding of the immune responses against current and future pandemics and will indicate a rational way to design antigens for effective vaccination strategies.

## Materials and Methods

### CD4 and B cell immunodominant epitopes selection and analysis

To focus the analysis on HLA-independent immunodominant determinants, experimentally validated CD4 and B epitopes that were found to be dominant in at least two different species / mouse strains or multiple patients were selected for further testing. B cell epitopes were arbitrarily classified as distant (>15aa), adjacent (<15aa) or overlapping (<0aa) with dominant CD4 epitopes based on their distance and positioning within the antigenic linear sequence. As such, peptides distant less than 15aa were considered likely to be near enough for the Ab to suppress the CD4 epitope.

### Crystal structures visualization

To highlight inter-positioning of B and CD4 dominant epitopes in 3D structures, above-mentioned PDB files were modified using Pymol (2.3.5). CD4 dominant epitopes were labeled in blue, dominant B epitopes in red and overlapping regions in magenta. Amino acid essential for bnAb binding to antigens are listed in indicated figures and those overlapping with dominant CD4 epitopes are highlighted in magenta.

### Statistical analysis

Prism software (GraphPad 9.0.1) was used for all statistical analysis. Chi-squared test was performed to evaluate the relationship between immunogenicity and the relative distribution of B and T cell epitopes within antigens. P<0.05 was considered significant. In summary graphs, points indicate samples and horizontal lines are means. Error bars indicated standard error mean (SEM).

## Figure legends

**Fig. S1.**
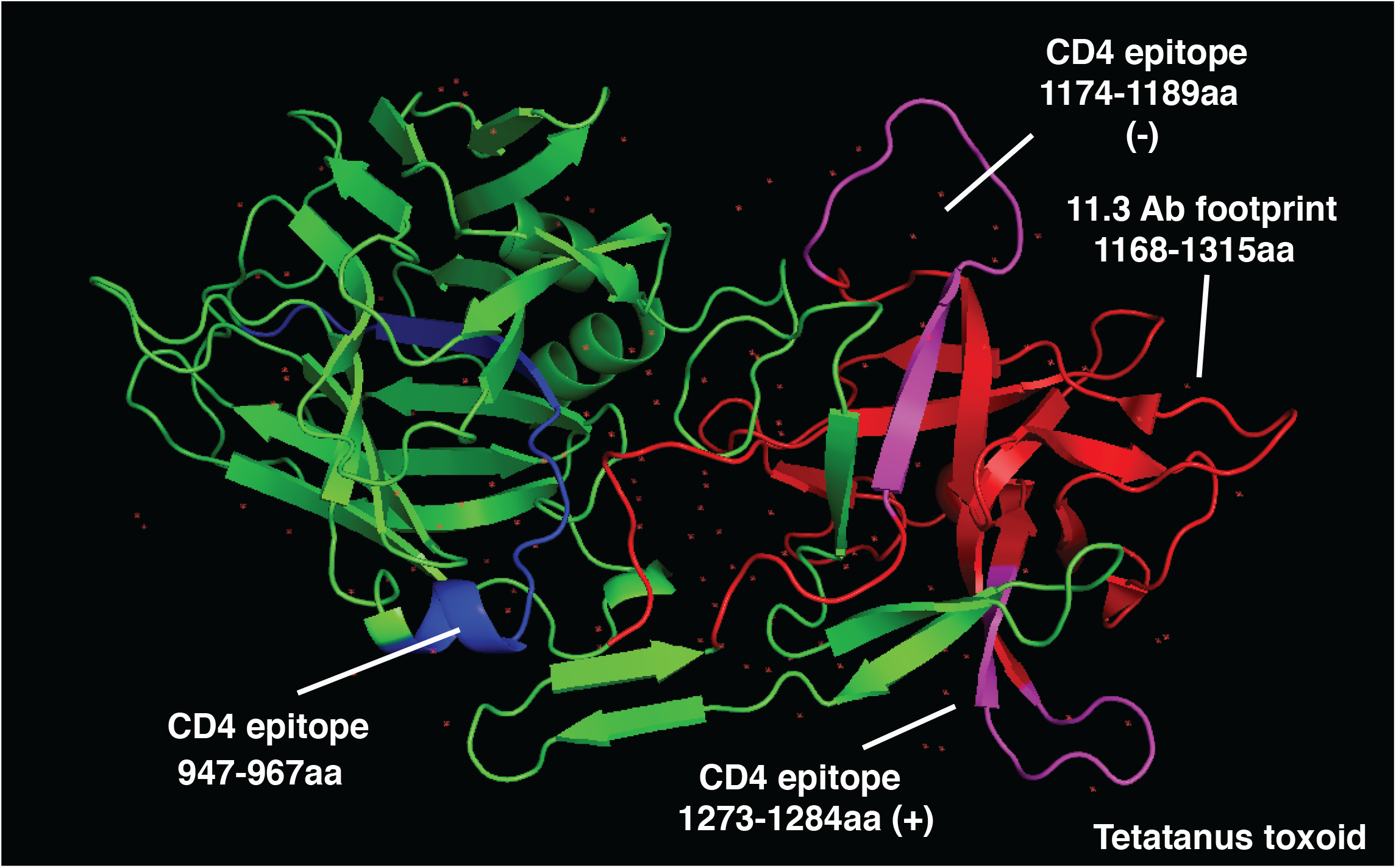
Crystal structure of tetanus toxoid (**PDB:1AF9**) is shown. Antibody footprint binding region is depicted in red, while CD4 determinants are highlighted in blue.

**Fig. S2.**
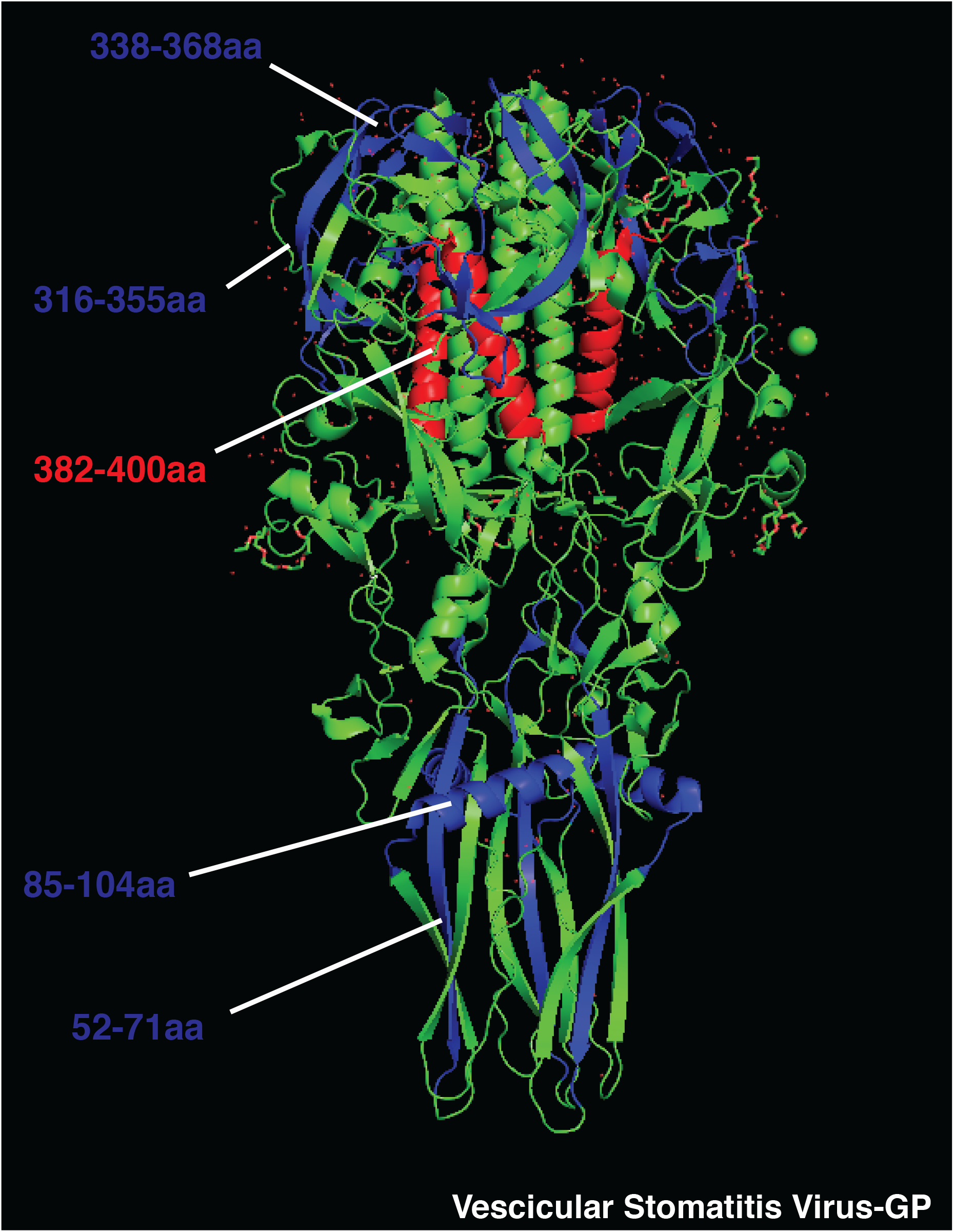
Crystal structure of Vescicular Stomatitis Virus glycoprotein (**PDB:512M**) is shown. Immunodominant B cell epitopes are depicted in red, while dominant CD4 determinants are highlighted in blue.

**Fig. S3.**
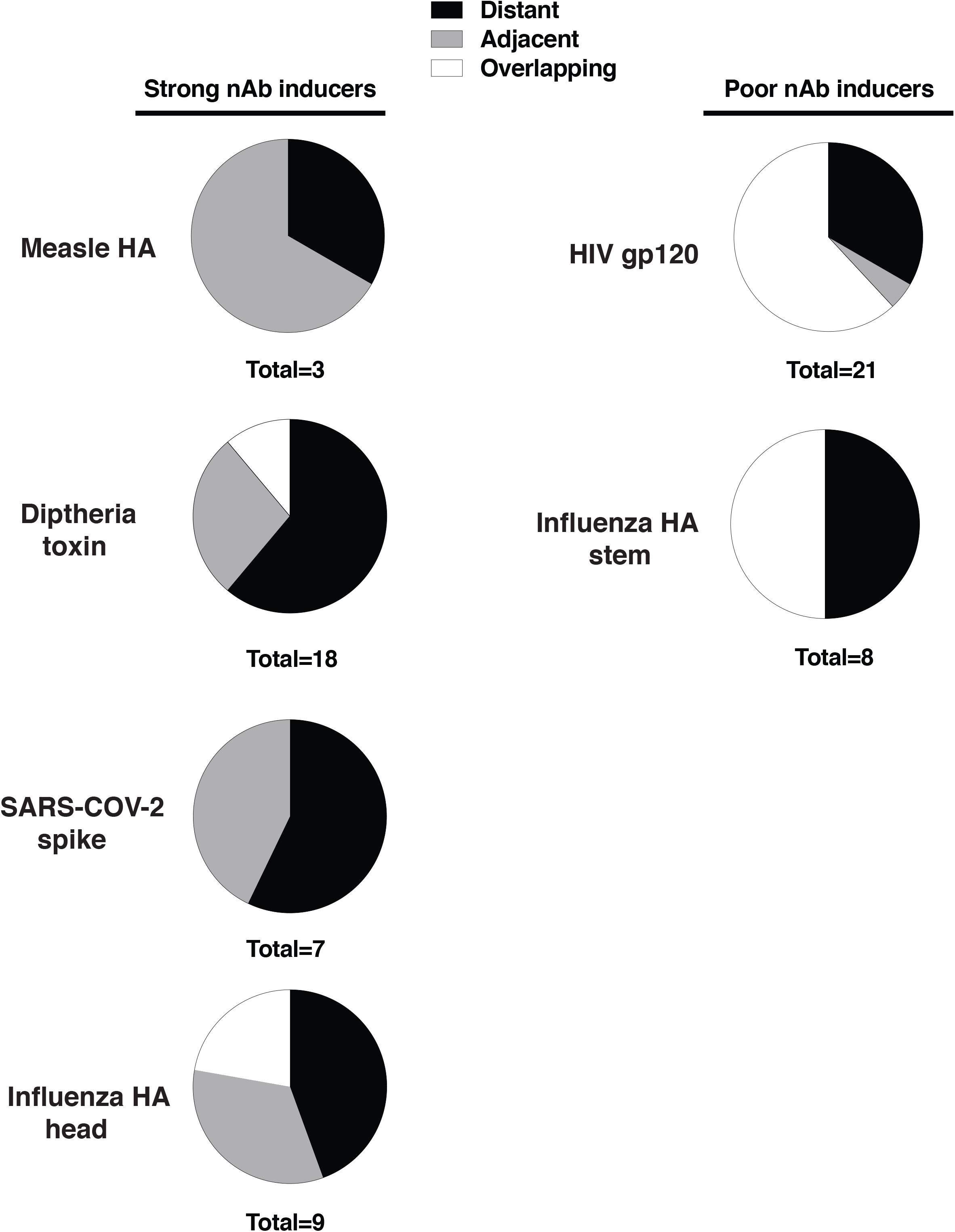
Pie charts showing percentages of distant (black), adjacent (gray) or overlapping (white) B epitopes in indicated antigens (good inducers, left; poor inducers right).

## Author contributions

RB and MDG conceived the idea, processed the literature, performed the analysis and wrote the paper. MDG prepared the figures.

## Acknowledgments

We thank prof. Jason G. Cyster and Dr. Giulio Cavalli for reading the manuscript and for providing insightful comments.

## References

1. Corti, D. & Lanzavecchia, A. Broadly neutralizing antiviral antibodies. Annual Review of Immunology 31, (2013).

2. Rees, A. R. Understanding the human antibody repertoire. MAbs 12, 1–16 (2020).

3. Abbott, R. K. & Crotty, S. Factors in B cell competition and immunodominance. Immunol. Rev. 296, 120–131 (2020).

4. Cyster, J. G. Germinal Centers: Gaining Strength from the Dark Side. Immunity 43, 1026–1028 (2015).

5. Bannard, O. & Cyster, J. G. Germinal centers: programmed for affinity maturation and antibody diversification. Current Opinion in Immunology 45, 21–30 (2017).

6. Cyster, J. G. Shining a light on germinal center B cells. Cell 143, 503–505 (2010).

7. Okada, T. & Cyster, J. G. B cell migration and interactions in the early phase of antibody responses. Current Opinion in Immunology 18, 278–285 (2006).

8. De Giovanni, M. & Iannacone, M. In vivo imaging of adaptive immune responses to viruses. Curr. Opin. Virol. 28, 102–107 (2018).

9. Cyster, J. G. B cell follicles and antigen encounters of the third kind. Nature Immunology 11, 989–996 (2010).

10. Paige, C. J. & Wu, G. E. The B cell repertoire. FASEB J. 3, 1818–1824 (1989).

11. Dunn-Walters, D. K. & Ademokun, A. A. B cell repertoire and ageing. Current Opinion in Immunology 22, 514–520 (2010).

12. Angeletti, D. et al. Defining B cell immunodominance to viruses. Nat. Immunol. 18, 456–463 (2017).

13. Barouch, D. H. Challenges in the development of an HIV-1 vaccine. Nature 455, 613–619 (2008).

14. Myszka, D. G. et al. Energetics of the HIV gp120-CD4 binding reaction. Proc. Natl. Acad. Sci. U. S. A. 97, 9026–9031 (2000).

15. Broecker, F. et al. Immunodominance of Antigenic Site B in the Hemagglutinin of the Current H3N2 Influenza Virus in Humans and Mice. J. Virol. 92, 1–13 (2018).

16. Akram, A. & Inman, R. D. Immunodominance: A pivotal principle in host response to viral infections. Clin. Immunol. 143, 99–115 (2012).

17. Cassotta, A. et al. Deciphering and predicting CD4+ T cell immunodominance of influenza virus hemagglutinin. J. Exp. Med. 217, (2020).

18. Galkin, A. et al. HIV-1 gp120–CD4-Induced Antibody Complex Elicits CD4 Binding Site–Specific Antibody Response in Mice. J. Immunol. 204, 1543–1561 (2020).

19. Jardine, J. G. et al. HIV-1 broadly neutralizing antibody precursor B cells revealed by germline-targeting immunogen. Science (80-.). 351, 1458–1463 (2016).

20. Chen, Y. et al. Influence of HLA alleles on the rate of progression of vertically transmitted HIV infection in children: Association of several HLA-DR13 alleles with long-term survivorship and the potential association of HLA-A*2301 with rapid progression to AIDS. Hum. Immunol. 55, 154–162 (1997).

21. Malhotra, U. et al. Role for HLA class II molecules in HIV-1 suppression and cellular immunity following antiretroviral treatment. J. Clin. Invest. 107, 505–517 (2001).

22. Kingdom, U. & Heights, A. Suppressive Effect of Antibody on Processing of T Cell Epitopes By Colin Watts* and Antonio Lanzavecchiar. 178, (1993).

23. Antonio Lanzavecchia. Antigen-specfic interaction between T and B cells. Nature 314, 537–539 (1985).

24. Lanzavecchia, A. Receptor-mediated antigen uptake and its effect on antigen presentation to class II-restricted T lymphocytes. Annu. Rev. Immunol. 8, 773–793 (1990).

25. Sanjuán, R., Nebot, M. R., Peris, J. B. & Alcamí, J. Immune Activation Promotes Evolutionary Conservation of T-Cell Epitopes in HIV-1. PLoS Biol. 11, (2013).

26. Levitz, L. et al. Conservation of HIV-1 T cell epitopes across time and clades: Validation of immunogenic HLA-A2 epitopes selected for the GAIA HIV vaccine. Vaccine 30, 7547–7560 (2012).

27. Lu, C. L. et al. Enhanced clearance of HIV-1-infected cells by broadly neutralizing antibodies against HIV-1 in vivo. Science (80-.). 352, 1001–1004 (2016).

28. Paules, C. I. et al. The hemagglutinin A stem antibody MEDI8852 prevents and controls disease and limits transmission of pandemic influenza viruses. J. Infect. Dis. 216, 356–365 (2017).

29. Granger, B. L. Accessibility and contribution to glucan masking of natural and genetically tagged versions of yeast wall protein 1 of Candida albicans. PLoS One 13, (2018).

30. Harris, A. K. et al. Structure and accessibility of HA trimers on intact 2009 H1N1 pandemic influenza virus to stem region-specific neutralizing antibodies. Proc. Natl. Acad. Sci. U. S. A. 110, 4592–4597 (2013).

31. Lusso, P. et al. Cryptic Nature of a Conserved, CD4-Inducible V3 Loop Neutralization Epitope in the Native Envelope Glycoprotein Oligomer of CCR5-Restricted, but Not CXCR4-Using, Primary Human Immunodeficiency Virus Type 1 Strains. J. Virol. 79, 6957–6968 (2005).

32. Mohabatkar, H. & Kar, S. K. Prediction of exposed domains of envelope glycoprotein in Indian HIV-1 isolates and experimental confirmation of their immunogenicity in humans. Brazilian J. Med. Biol. Res. 37, 675–681 (2004).

33. Xiao, X., Chen, W., Feng, Y. & Dimitrov, D. S. Maturation pathways of cross-reactive HIV-1 neutralizing antibodies. Viruses 1, 802–817 (2009).

34. Li, Y. et al. Mechanism of Neutralization by the Broadly Neutralizing HIV-1 Monoclonal Antibody VRC01. J. Virol. 85, 8954–8967 (2011).

35. Dreja, H., Pade, C., Chen, L. & McKnight, Á. CD4 binding site broadly neutralizing antibody selection of HIV-1 escape mutants. J. Gen. Virol. 96, 1899–1905 (2015).

36. Sun, M., Li, Y., Zheng, H. & Shao, Y. Recent progress toward engineering HIV-1-specific neutralizing monoclonal antibodies. Front. Immunol. 7, (2016).

37. Wei, C. J. et al. Next-generation influenza vaccines: opportunities and challenges. Nature Reviews Drug Discovery 19, 239–252 (2020).

38. Yuan, M. et al. A highly conserved cryptic epitope in the receptor binding domains of SARS-CoV-2 and SARS-CoV. Science (80-.). 368, 630–633 (2020).

39. Murin, C. D. et al. Structural Basis of Pan-Ebolavirus Neutralization by an Antibody Targeting the Glycoprotein Fusion Loop. Cell Rep. 24, 2723–2732.e4 (2018).

40. Mitran, C. J. et al. Antibodies to cryptic epitopes in distant homologues underpin a mechanism of heterologous immunity between plasmodium vivax PvDBP and plasmodium falciparum VAR2CSA. MBio 10, (2019).

41. Pappas, L. et al. Rapid development of broadly influenza neutralizing antibodies through redundant mutations. Nature 516, 418–422 (2014).

42. Cohen, O. J., Kinter, A. & Fauci, A. S. Host factors in the pathogenesis of HIV disease. Immunological Reviews 159, 31–48 (1997).

43. Keet, I. P. M. et al. Consistent associations of HLA class I and II and transporter gene products with progression of human immunodeficiency virus type 1 infection in homosexual men. J. Infect. Dis. 180, 299–309 (1999).

44. Kaslow, R.. et al. A1, Cw7, B8, DR3 HLA antigen combination associated with rapid decline of T-helper lymphocytes in HIV-1 infection. Lancet 335, 927–930 (1990).

45. Landais, E. et al. Broadly Neutralizing Antibody Responses in a Large Longitudinal Sub-Saharan HIV Primary Infection Cohort. PLoS Pathog. 12, (2016).

46. Roche, P. A. & Furuta, K. The ins and outs of MHC class II-mediated antigen processing and presentation. Nat. Rev. Immunol. 15, 203–216 (2015).

47. Casten, L. A. & Pierce, S. K. Receptor-mediated B cell antigen processing. Increased antigenicity of a globular protein covalently coupled to antibodies specific for B cell surface structures. J. Immunol. 140, 404–40410 (1988).

48. Melchers, F. & Messer, W. Enhanced stability against heat denaturation of E. coli wild type and mutant β-galactosidase in the presence of specific antibodies. Biochem. Biophys. Res. Commun. 40, 570–575 (1970).

49. Jemmerson, R. & Paterson, Y. Mapping epitopes on a protein antigen by the proteolysis of antigen-antibody complexes. Science (80-.). 232, 1001–1004 (1986).

50. Davidson, H. W. & Watts, C. Epitope-directed processing of specific antigen by B lymphocytes. J. Cell Biol. 109, 85–92 (1989).

51. Berzofsky, J. A. An Ia-restricted epitope-specific circuit regulating T cell-B cell interaction and antibody specificity. Surv. Immunol. Res. 2, 223–229 (1983).

52. Manca, F. et al. Constraints in TtJB cooperation related to epitope topology on E. coli βtJgalactosidase. I. The fine specificity of T cells dictates the fine specificity of antibodies directed to conformationtJ dependent determinants. Eur. J. Immunol. 15, 345–350 (1985).

53. Manca, F. et al. Differential activation of T cell clones stimulated by macrophages exposed to antigen complexed with monoclonal antibodies. A possible influence of paratope specificity on the mode of antigen processing. J. Immunol. 140, 2893–2898 (1988).

54. Ozaki, S. & Berzofsky, J. A. Antibody conjugates mimic specific B cell presentation of antigen: Relationship between T and B cell specificity. J. Immunol. 138, 4133–4142 (1987).

55. Celada, F. & Sercarz, E. E. Preferential pairing of T-B specificities in the same antigen: the concept of directional help. Vaccine 6, 94–98 (1988).

56. Simitsek, P. D., Campbell, D. G., Lanzavecchia, A., Fairweather, N. & Watts, C. Modulation of antigen processing by bound antibodies can boost or suppress class II major histocompatibility complex presentation of different T cell determinants. J. Exp. Med. 181, 1957–1963 (1995).

57. Lanzavecchia, A. Antigen presentation by B lymphocytes: a critical step in T-B collaboration. Curr. Top. Microbiol. Immunol. 130, 65–78 (1986).

58. Marttila, J., Ilonen, J., Norrby, E. & Salmi, A. Characterization of T cell epitopes in measles virus nucleoprotein. J. Gen. Virol. 80, 1609–1615 (1999).

59. Tahara, M. et al. Functional and Structural Characterization of Neutralizing Epitopes of Measles Virus Hemagglutinin Protein. J. Virol. 87, 666–675 (2013).

60. Diethelm-Okita, B. M., Okita, D. K., Banaszak, L. & Conti-Fine, B. M. Universal epitopes for human CD4+ cells on tetanus and diphtheria toxins. J. Infect. Dis. 181, 1001–1009 (2000).

61. Zhu, S. et al. Hydrogen-Deuterium Exchange Epitope Mapping Reveals Distinct Neutralizing Mechanisms for Two Monoclonal Antibodies against Diphtheria Toxin. Biochemistry 58, 646–656 (2019).

62. De-Simone, S. G. et al. Epitope mapping of the diphtheria toxin and development of an ELISA-specific diagnostic assay. Vaccines 9, (2021).

63. Keil, W. & Wagner, R. R. Epitope mapping by deletion mutants and chimeras of two vesicular stomatitis virus glycoprotein genes expressed by a vaccinia virus vector. Virology 170, 392–407 (1989).

64. Burkhart, C. et al. Characterization of T-helper epitopes of the glycoprotein of vesicular stomatitis virus. J. Virol. 68, 1573–1580 (1994).

65. Hu, B., Guo, H., Zhou, P. & Shi, Z. L. Characteristics of SARS-CoV-2 and COVID-19. Nat. Rev. Microbiol. 19, 141–154 (2021).

66. Pinto, D. et al. Broad betacoronavirus neutralization by a stem helix–specific human antibody. Science (80-.). 373, 1109–1116 (2021).

67. Pinto, D. et al. Cross-neutralization of SARS-CoV-2 by a human monoclonal SARS-CoV antibody. Nature 583, 290–295 (2020).

68. Lu, S. et al. The immunodominant and neutralization linear epitopes for SARS-CoV-2 National Key Laboratory of Biochemical Engineering, Institute of Process Engineering, Chinese Academy of Sciences, Beijing 100190, China Innovation Academy for Green Manufacture, Ch. (2020).

69. Poh, C. M. et al. Two linear epitopes on the SARS-CoV-2 spike protein that elicit neutralising antibodies in COVID-19 patients. Nat. Commun. 11, (2020).

70. Peng, Y. et al. Broad and strong memory CD4+ and CD8+ T cells induced by SARS-CoV-2 in UK convalescent individuals following COVID-19. Nat. Immunol. 21, 1336–1345 (2020).

71. Greaney, A. J. et al. Mapping mutations to the SARS-CoV-2 RBD that escape binding by different classes of antibodies. Nat. Commun. 12, (2021).

72. Bertoglio, F. et al. A SARS-CoV-2 neutralizing antibody selected from COVID-19 patients binds to the ACE2-RBD interface and is tolerant to most known RBD mutations. Cell Rep. 36, (2021).

73. Dai, G., Steede, N. K. & Landry, S. J. Allocation of Helper T-cell Epitope Immunodominance According to Three-dimensional Structure in the Human Immunodeficiency Virus Type I Envelope Glycoprotein gp120. J. Biol. Chem. 276, 41913–41920 (2001).

74. Landry, S. J. Helper T-cell epitope immunodominance associated with structurally stable segments of hen egg lysozyme and HIV gp120. J. Theor. Biol. 203, 189–201 (2000).

75. Schrier, R. D. et al. T cell recognition of HIV synthetic peptides in a natural infection. J. Immunol. 142, 1166–1176 (1989).

76. Surman, S. et al. Localization of CD4 L T cell epitope hotspots to exposed strands of HIV envelope glycoprotein suggests structural influences on antigen processing. 98, 4587–4592 (2001).

77. Wahren, B. et al. HIV-1 peptides induce a proliferative response in lymphocytes from infected persons. J. Acquir. Immune Defic. Syndr. 2, 448–456 (1989).

78. Hioe, C. E. et al. Inhibition of Human Immunodeficiency Virus Type 1 gp120 Presentation to CD4 T Cells by Antibodies Specific for the CD4 Binding Domain of gp120. J. Virol. 75, 10950–10957 (2001).

79. Tuen, M. et al. Characterization of antibodies that inhibit HIV gp120 antigen processing and presentation. Eur. J. Immunol. 35, 2541–2551 (2005).

80. Lee, J. H. et al. Modulating the quantity of HIV Env-specific CD4 T cell help promotes rare B cell responses in germinal centers. J. Exp. Med. 218, (2020).

81. Landry, S. J. Three-Dimensional Structure Determines the Pattern of CD4+ T-Cell Epitope Dominance in Influenza Virus Hemagglutinin. J. Virol. 82, 1238–1248 (2008).

